# In vitro and in vivo comparison of the MRI glucoCEST properties between native glucose and 3OMG in a murine tumor model

**DOI:** 10.1101/2021.03.15.435387

**Authors:** Annasofia Anemone, Martina Capozza, Francesca Arena, Sara Zullino, Paola Bardini, Enzo Terreno, Dario Livio Longo, Silvio Aime

## Abstract

**Purpose:** D-Glucose and 3-O-Methyl-D-glucose (3OMG) have been shown to provide contrast in MRI-CEST images. However, a systematic comparison between these two molecules has not yet been performed. This study dealt with the assessment of the effect of pH, saturation power level (B_1_) and magnetic field strength (B_0_) on the MRI-CEST contrast with the aim of comparing the *in vivo* CEST contrast detectability of these two agents in the glucoCEST procedure.

**Methods:** Phosphate buffered solutions of D-Glucose or 3OMG (20 mM) were prepared at different pH values and Z-spectra acquired at several B_1_ levels and at 37°C. *In vivo* glucoCEST images were obtained at 3 T and 7 T over a period of 30 min after injection of D-Glucose or 3OMG (at the doses of 1.5 and 3 g/kg) in a murine melanoma tumour model.

**Results:** A markedly different pH dependence of CEST response was observed in vitro for D-Glucose and 3OMG. The glucoCEST contrast enhancement in the tumour region following the intravenous administration (at the dose 3 g/kg) resulted to be comparable for both the molecules: 1-2% at 3 T and 2-3% at 7 T. The ST% resulted almost constant for 3OMG over the 30 min period, whereas a significant increase in the case of D-Glucose was detected.

**Conclusion:** Our results show similar CEST contrast efficiency but different temporal kinetics for the metabolizable and the non-metabolizable glucose derivatives in tumour murine models when administered at the same doses.

## 1. Introduction

*In vivo* imaging techniques are currently used to detect tumour and to monitor the response to therapy. Magnetic Resonance Imaging (MRI) often makes use of contrast agents to augment physiological information to the anatomical resolution of its images ^1^. Nowadays, much attention is devoted to the characterization of tumour metabolism as it is recognised that the knowledge of the metabolic state of cancer cells is a crucial information in the diagnostic assessment of the disease. Glucose, being the primary source of energy, is of course under intense scrutiny as a metabolic tracer ^2^. Positron emission tomography (PET) exploits the increased glucose uptake from tumour cells to report the accumulation of 2-Deoxy-2-[^18^F]-fluoroglucose ([^18^F]-FDG), a radioactive glucose analogue, and to extract precious information on the ongoing metabolism of the cancer cells ^3^. Indeed, this method is daily used in clinical settings to image primary and metastatic tumours. Major issues that hamper the PET modality are associated with the use of radioactive compounds and to the radiation doses that the patients receive when the PET is carried out in association with computed tomography (CT) to provide the required anatomical resolution. These issues make its application not suitable for all patients (e.g. pregnant women and pediatric application are commonly excluded from PET studies) and quite expensive ^4^.

Over the past decade, MRI-CEST (Chemical Exchange Saturation Transfer) methods have attracted interest as non-invasive alternatives to study tumour metabolism and its microenvironment ^5–7^; CEST endogenous contrast has been widely explored in pre-clinical and clinical studies ^8^. Magnetization transfer between exchangeable protons from amine ^9,10^, amide ^11,12^ or hydroxyl group ^13,14^ resonating between 1 ppm and 3 ppm downfield from the water resonance can be exploited for CEST applications. Much attention has been devoted to the use of glucose as MRI-CEST agent as it would yield a significant cost reduction and an increase in safety and accessibility in comparison to PET, yet preserving and potentially improving its specificity for tumour characterization and evaluation of response to therapy.

Exploiting OH exchangeable protons as the source of the MRI CEST effect, natural glucose proved its usefulness in cancer detection. A first demonstration was provided when glucoCEST showed enhanced contrast in two human breast cancer cell lines orthotopically implanted in mice in agreement with FDG-PET enhancement ^15^. Later, on human colorectal tumour mouse xenograft model the glucoCEST signal compared to [^18^F]-FDG autoradiography provided further support to the view that glucoCEST is specific and a sensitive measure as much as FDG uptake ^16^. Moreover, dynamic glucose enhanced MRI was used to study malignant brain tumour and blood barrier breakdown also at clinical level ^17,18^. It was also shown that the sensitivity in glucose mapping concentration in brain could be increased by assessing the OH proton exchange by means of spin-lock MRI ^19,20^. Recently, in a detailed study of the exchange rate constant of glucose hydroxyl groups, one may access to optimal parameters for the in vivo glucoCEST/Chemical exchange-sensitive spin-lock (CESL) detection at clinical and ultra-high field strengths ^21–23^.

However, when compared to [^18^F]-FDG, glucose is rapidly metabolized through glycolysis, thus causing uncertainty about its concentration in tumour cells with a consequent decrease of the CEST signal. For this reason, glucose analogues that are not metabolized upon their uptake into tumour cells, are being considered as possible alternatives for MRI-CEST procedures. For instance, analogues such as 2-deoxy-D-glucose (2DG), dextran, sucralose, sucrose, glucosamine (GlcN) and N-acetyl-glucosamine (GlcNAc) that are phosphorylated as glucose ^24–28^ and analogues that do not undergo phosphorylation such as 2-O-Methyl-D-glucose (2OMG), 3-O-Methyl-D-glucose (3OMG) and 6-deoxy-D-glucose (6DG) are under intense study ^29–31^. Among the first group, GlcN is considered worth of note for its excellent safety profile ^28^ and for the presence in its structure of the amino peak that yields a CEST signal that is more shifted than the hydroxyl ones from water, thus making the CEST response more efficient, in particular at the clinical fields. In the second group, 3-O-Methyl-D-glucose (3OMG) appears an interesting alternative as it enters the cells through the glucose transporters (GLUT-1 and 3). 3OMG displayed higher and longer lasting CEST signal compared to D-glucose for the same type of murine tumour ^32^, but detailed studies of its toxicology are still lacking, even if it did not induce any physiological or behavioural changes for various dosages in mice and rats ^33^.

Although successful proofs of concept have been provided and clinical investigations have been reported for D-glucose ^20,34,35^, to date no comprehensive studies investigating D-glucose and 3OMG on the same tumour model and under the same experimental conditions are available. The aim of this work is to evaluate systematically the effects of pH, saturation power level (B_1_) and magnetic field strength (B_0_) on the generation of CEST contrast of D-glucose and 3OMG for a proper comparison between the two molecules. Moreover, we assessed their *in vivo* capability to provide contrast at two field strengths (3 T and 7T) when administered to a murine melanoma model upon an intravenous injection at two different doses (1.5 and 3 g/kg).

## 2. Methods

### 2.1 Chemicals

D-glucose and 3-O-methyl-D-glucose powder for *in vitro* studies were obtained from Sigma-Aldrich (Milan, Italy). Solutions of D-glucose and 3OMG for *in vitro* studies were prepared in 10 mM phosphate-buffered saline (PBS 1X). Glucose or 3-O-methyl-D-glucose injectable solution for *in vivo* studies was prepared dissolving the powder in saline solution to obtain a 3 M solution at neutral pH (pH 7.4). The solution was then filtered with a 200 nm membrane filters to preserve the suspensions from bacterial contamination. 3OMG powder for *in vivo* studies was kindly provided by Almac (Almac, UK).

### 2.2 Phantom preparation

Phantoms containing different vials of 10mM phosphate-buffered solutions (PBS 1X) of D-Glucose and 3OMG were prepared starting from a 20 mM solution. Each solution was then titrated to reach the intended pH of 7.4, 7.0, 6.8, 6.6, 6.4, 6.2, and 6.0, respectively.

### 2.3 Tumour animal model

#### 2.3.1 Cell culture

B16-F10 (mouse melanoma cells) were obtained from American Type Culture Collection (ATCC). B16-F10 cells were cultured in EMEM supplemented with 10% FBS, 100 μg/mL penicillin and 100 μg/mL streptomycin. The cells were grown at 37°C in a humidified atmosphere containing 5% CO_2_. At confluence, cells were detached by adding 1 mL of Trypsin-EDTA solution (0.25 %_w/v_ Trypsin, 0.53 mM EDTA). EMEM, FBS and Trypsin were purchased from Lonza (Lonza Sales AG, Verviers, Belgium). The penicillinstreptomycin mixture was purchased from Sigma Chemical Co., St. Louis, MO, USA.

#### 2.3.2 Subcutaneous tumour cells implantation in mice

Male 8 weeks old C57BL/6 mice (Charles River Laboratories Italia S.r.l., Calco, Italia) were inoculated with 5.0 × 10^5^ B16-F10 melanoma cells in both flanks 10 days before imaging. Mice were maintained under specific pathogen free conditions in the animal facility of the Centre for Preclinical Imaging, University of Turin, and treated in accordance with the University Ethical Committee and European guidelines under directive 2010/63. B16-F10 tumour bearing mice were randomly divided in 8 groups (3 to 5 mice each) to investigate the elicited CEST contrast after intravenous injection of D-Glucose or 3OMG at the two doses of 1.5 g/kg and 3 g/kg, at two main magnetic fields of 3 T and 7 T, respectively.

Before imaging, mice were anesthetized with isoflurane and placed on the MRI bed and monitored through an air pillow located below the animal (SA Instruments, Stony Brook, NY; USA). The tail vein was cannulated with a catheter with a 27-gauge needle.

### 2.4 MRI-CEST protocol and analysis

*In vitro* Z-spectra were acquired on a Bruker BioSpec (3 T) scanner equipped with a 1H quadrature coil and on a Bruker Avance 300 (7 T) scanner equipped with a micro 2.5 imaging probe. The experiments were carried out at 37°C by irradiating the sample with a single continuous wave pre-saturation block pulse (1.0, 2.0 and 3.0 μT) applied for 5 sec. The saturation frequency offset was varied between 10 and −10 ppm with a frequency resolution of 0.1 ppm. MR images were acquired using a Spin Echo RARE sequence (TR/TE/NEX/Rare Factor 10.0 sec/5.4 msec/2/64) with centric encoding, field of view = 3 × 3 cm; slice thickness = 2 mm; matrix =64 × 64.

*In vivo* studies were carried out on a Bruker Pharmascan (7 T) scanner (Bruker Biospin, Ettlingen, Germany) equipped with a 30 mm 1H quadrature coil and on a Bruker BioSpec (3 T) scanner equipped with a 30 mm 1H quadrature coil. After the scout image, T_2w_ anatomical images were acquired with a RARE sequence and the same geometry was used for the following CEST experiments. The glucoCEST (before and after injection of D-Glucose or 3OMG) images were obtained by irradiating the animal with a single continuous wave presaturation block pulse of 2 μT applied for 5 sec. Z-spectra were sampled with 61 frequency offsets in the range ±10 ppm with a step size of 0.2 ppm in the range ±6 ppm. CEST images were recorded with a singleshot Fast Spin Echo sequence with centric encoding (TR 6.0 s, TE 4.7 ms, Rare Factor 64, field of view = 3 cm × 3 cm; slice thickness = 2 mm; matrix = 64 × 64). CEST images were acquired before and every 6 min following the injection up to 32 min.

In-house MATLAB scripts (The Mathworks, Inc., Natick, MA, USA) were used to process all the CEST images. Firstly, anatomical and Z-spectrum images were segmented by using an intensity-threshold filter; secondly, Z-spectra were interpolated on a voxel-by-voxel basis by smoothing splines ^36^ to identify the correct position of the bulk water and remove artifacts arising from B_0_ inhomogeneity. Then, the interpolated Z-spectrum was shifted to make bulk water resonance match to the zero frequency and corrected intravoxel saturation transfer (ST) effects were calculated with asymmetry analysis ^37^. To remove CEST effect arising from noisy data, a second filter was applied by calculating for the interpolating curve the coefficient of determination (R^2^) and to consider the signal-to-noise ratio of single voxels (noisy Z-spectra present low R^2^ values); in the ST% calculation only voxels with high R^2^ (>0.97) were considered. The ST effect for glucose or 3OMG was estimated from the expression:

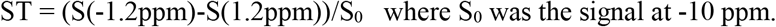

Results are reported as:

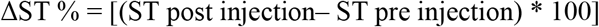

The fraction of enhanced pixel reports on the percentage of pixels showing a ΔST% greater than zero in the manually defined tumour region of interest (ROI).

### 2.5 Statistical analysis

GraphPad Prism 7 software (GraphPad Inc, San Diego, California, USA) was used for statistical analysis. Data are presented as mean ± SD unless otherwise stated. One-way ANOVA analysis and Dunnet’s multiple comparison test were used to test for statistically significant differences between the ΔST measurements along time. For all tests, a P value < 0.05 was considered statistically significant.

## 3. Results

### 3.1 In vitro MRI-CEST Characterization

To assess the magnetic field and pH dependent properties of D-Glucose and 3-O-methyl-D-glucose, phantoms containing the solutions at different pH values were investigated. Z-spectra were acquired on two MRI scanners, namely on a high field 7 T scanner and on a preclinical scanner working at clinical field of 3 T, respectively. The chemical shift (from water resonance) of 0.8 ppm for D-glucose and of 1.2 ppm for 3-O-methyl-D-glucose were chosen as they correspond to the highest CEST signals. The saturation transfer effect at 7 T measured at 37°C for 20 mM D-Glucose solution at different pH values using 3.0 μT of saturation pulse, is shown in Figure 1. The CEST effects calculated from the asymmetry analysis in the B_0_-corrected Z-spectra generated by D-Glucose reached 36% between pH 6.4 and 6.0 (Figure 1A and C). The CEST effect appears markedly pH dependent as at neutral pH a net decrease in the saturation transfer effect is clearly observed. Figure 1B shows the B_0_-corrected Z-spectra generated by solutions containing 20 mM of 3OMG obtained by using the same acquisition parameters reported above for D-Glucose containing solutions. Compared to D-Glucose, the 3OMG signal reached higher CEST effect at neutral pH (around 35% between pH 7.0 and 6.8) and decreases at acidic pH values (25 % of ST effect at pH 6.0, figure 1D). At 3 T, broader Z-spectra and lower contrasts were detected (Figure 2 A and B); the highest ST effect for D-Glucose (close to 25%) was observed between pH 6.0 and 6.4 (Figure 2C), as detected in the high field experiment. 3OMG, under the same acquisition conditions, showed lower ST effects (close to 10 %) compared to D-Glucose between pH 6.0 and 6.4 (Figure 2D), whereas the highest effect (20 %) was detected at neutral pH 7.0.

**FIGURE 1.**
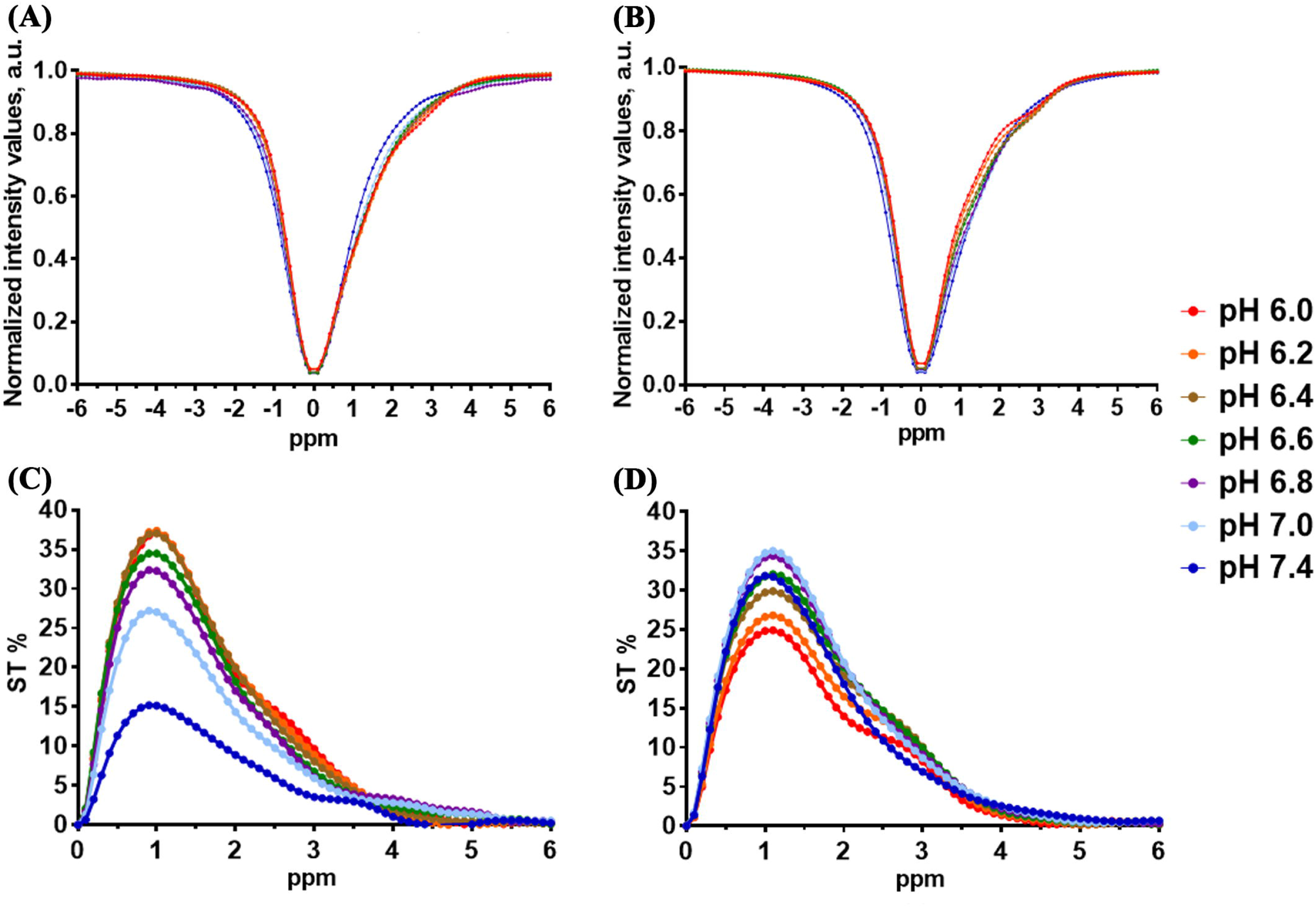
Z-Spectra and ST% effect plot of 20mM D-Glucose (A and C) and 3OMG (B and D) solution containing 10 mM of phosphate buffer as a function of pH values acquired at 37°C with a 7 T scanner (B_1_=3.0 μT).

**FIGURE 2.**
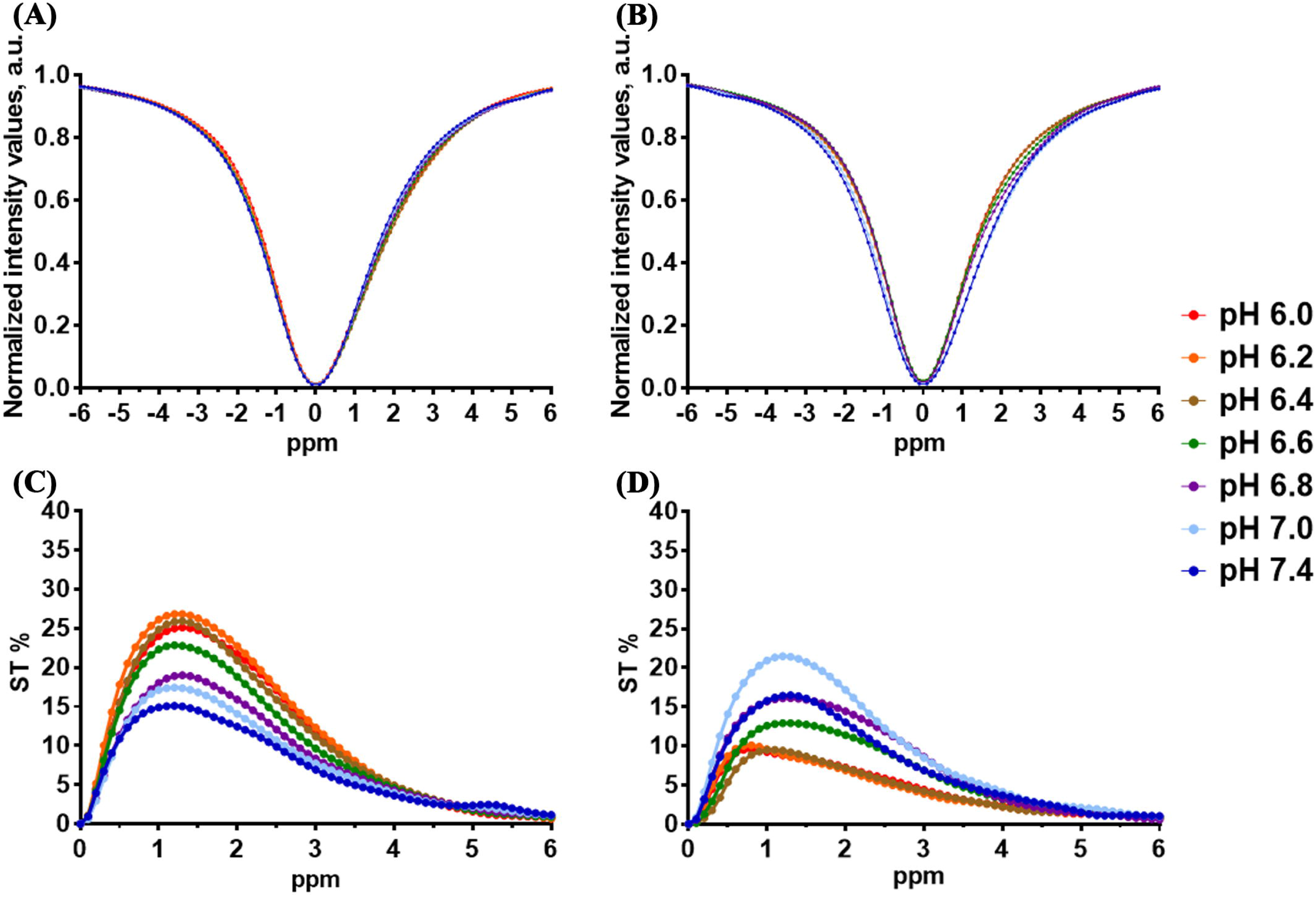
Z-Spectra and ST% effect plot of 20mM D-Glucose (A and C) and 3OMG (B and D) solution containing 10 mM of phosphate buffer as a function of pH values acquired at 37°C with a 3 T scanner (B_1_=3.0 μT).

Figure 3 reports the CEST contrast measured at 37 °C and 7 T or 3 T, at a frequency offset of 0.8 ppm for D-Glucose and 1.2 ppm for 3OMG. From these data one may get useful insights on the pH dependence for the two CEST agents at different saturation power values (1.0, 2.0 and 3.0 μT). For D-glucose an increase in the CEST effect around pH 6.0-6.2 at all the selected saturation power was observed (Figure 3A). The effect is more pronounced using 3.0 μT power pulse and decreases proportionally with increasing of pH, reaching the minimum at pH 7.4. In the case of 3OMG, an opposite trend compared to D-Glucose was observed (Figure 3B); the CEST response slightly increased starting from acidic pH up to pH 7.0, followed by a slight decrease at pH 7.4. At lower B_0_ field, the irradiation RF field of 3.0 μT resulted in the highest CEST contrast response for both Glucose and 3OMG. D-Glucose still displayed a higher CEST effect at acidic pH values, but the CEST contrast at 2.0 μT and at 3.0 μT were almost comparable (Figure 3C). For 3OMG the detected CEST effect showed a sharp increase from pH 6.4 that peaked at pH 7.0 and then decreased (Figure 3D).

**FIGURE 3.**
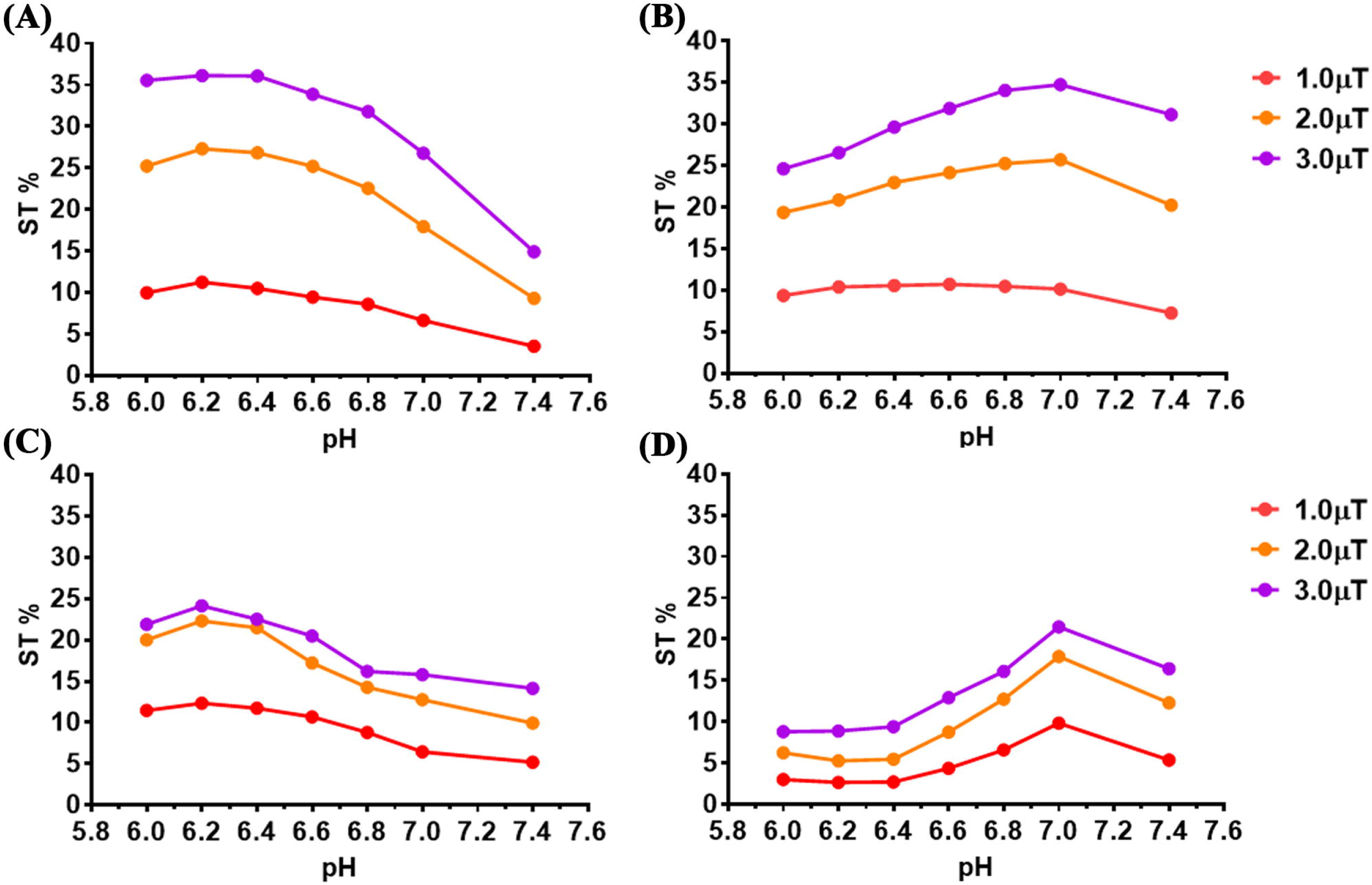
ST% effect plot of 20 mM D-Glucose (A and C) and 3OMG (B and D) solution containing 10 mM of phosphate buffer as a function of the RF saturation field (B_1_ =1.0, 2.0 and 3.0 μT) acquired by a 7 T (A and B) or by a 3 T MRI scanner (C and D) at 37°C.

### 3.2 In vivo CEST MRI studies

In order to determine the minimum dose needed to obtain sufficient glucoCEST contrast, mice inoculated with B16-F10 melanoma cells on both flanks underwent i.v. administration at 1.5 g/kg or 3 g/kg dose of D-Glucose or 3OMG, respectively. Pre- and post-contrast images were acquired with 7 T and 3 T preclinical scanners by applying a 2.0 μT RF pulse for 5 seconds. At 7 T, as shown in Figure 4A, the CEST contrast reached a value of 1.4 ± 0.2% at 6 min after a single bolus glucose i.v. injection of 1.5 g/kg. The CEST contrast slightly increased up to reach a value of 2.4 ± 0.7% 30 min after the injection. The calculated fraction of enhanced pixels (i.e. pixel showing a positive ΔST increase, Figure 4B) indicates a good coverage of the tumour region (from 66% right after the injection, to 78% after 30 min). A higher single D-Glucose dose of 3 g/kg improved the CEST signal over 30 min of observation, starting from a CEST contrast of 1.7 ± 0.3% that increased to 2.9 ± 0.7% (Figure 4C). Almost 80-85% of the tumour area showed a marked glucoCEST enhancement 12 min after the D-glucose injection (Figure 4D). ANOVA analysis indicated a significant difference in CEST contrast between 6 min from the injection and 12, 18, 24 or 30 min later (*P = 0.046, 0.022. 0.013 and 0.0016, respectively), at the highest dose and between 6 min and 30 min for the lower dose (*P = 0.0011).

**FIGURE 4.**
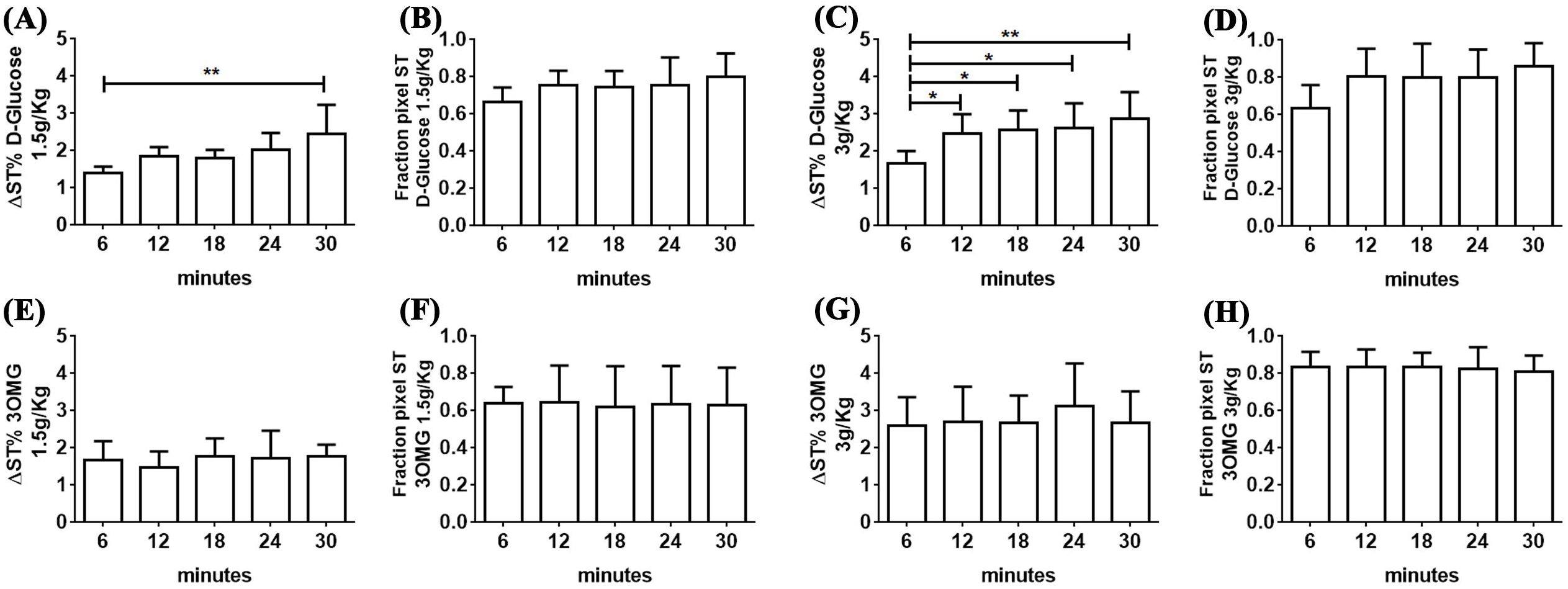
GlucoCEST contrast obtained at 7 T injecting D-Glucose at 1.5 g/kg (A) and 3 g/kg (C) dose or 3OMG at 1.5 g/kg (E) and 3 g/kg (G) dose via intravenous route and fraction of enhanced pixels at the corresponding injected dose (B, D for D-Glucose and F, H for 3OMG). Data are reported as the difference (ΔST %) between the ST effects before and after the intravenous injection.

Conversely, the 3OMG CEST contrast resulted stable over the 30 min observation time, but comparable in magnitude to that raised by D-glucose (Figure 4E, G). 3OMG at 1.5 g/kg dose showed a ΔST% of 1.7 ± 0.5% and a fraction of enhanced pixels of ca. 62-64% (Figures 4E and 4F). The 3 g/kg dose injection showed a marked increase of the CEST response (ΔST% 2.7 ± 0.9 %), with a fraction of enhanced pixels of 80-83% (Figure 4G and 4H). Representative parametric images for both D-glucose and 3OMG at the two doses and at the several time points (6-30 min after the i.v. administration) are shown in Figure 5.

**FIGURE 5.**
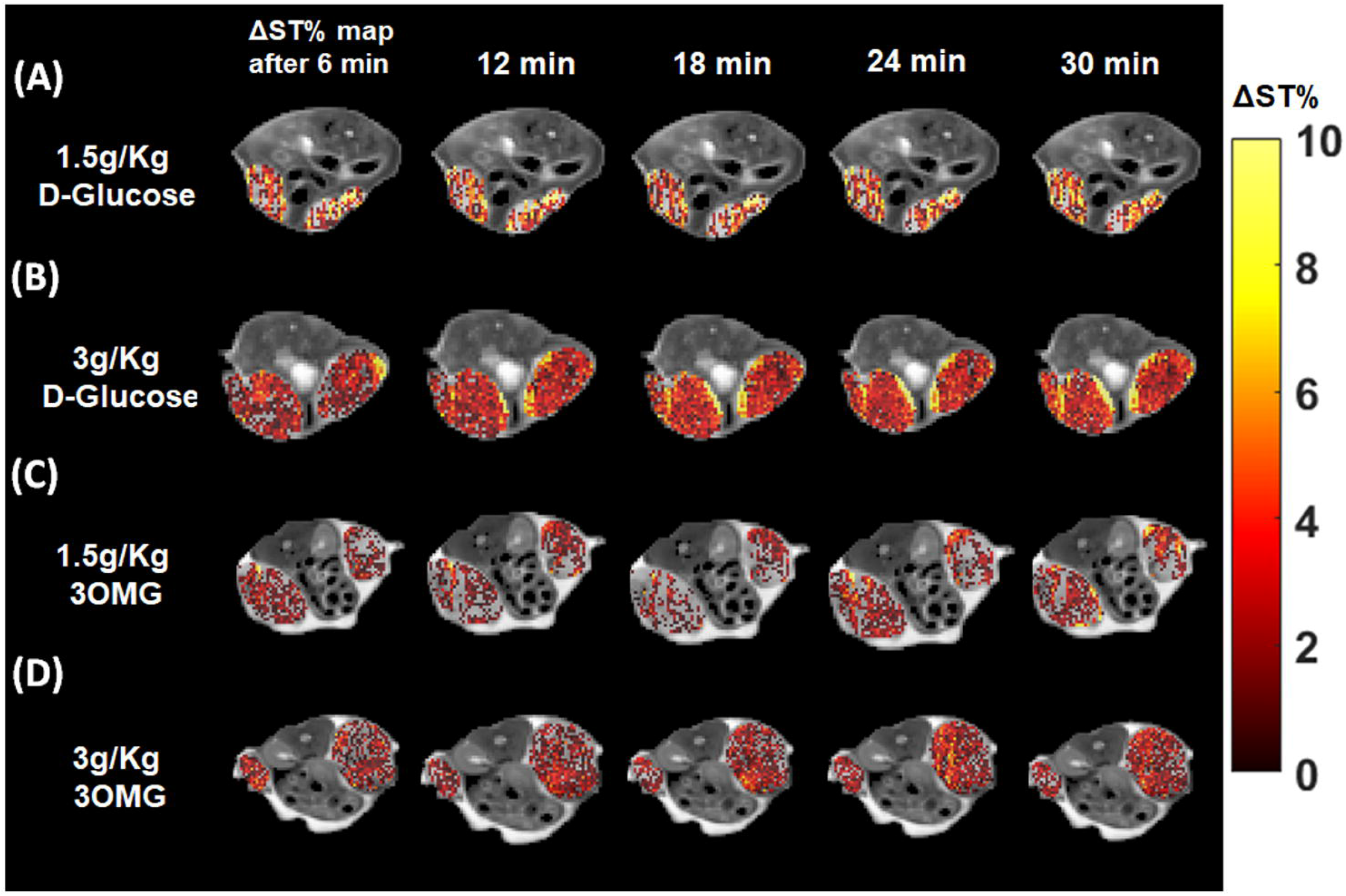
GlucoCEST ΔST% map obtained at 7 T injecting D-Glucose at 1.5 g/kg (A) and 3 g/kg (B) dose or 3OMG at 1.5 g/kg (C) and 3 g/kg (D) dose via intravenous route. Data are reported as the difference (ΔST %) between the ST effect before and after the intravenous injection, every 6 minutes from the injection. Parametric maps are overimposed to T_2w_ anatomical images and glucoCEST contrast is shown only in the tumour regions.

At 3 T, both D-Glucose and 3OMG displayed a smaller CEST contrast. The ΔST% of D-Glucose was 1.3 ± 0.3% from bolus injection of 1.5 g/kg (Figure 6A) and 1.5 ± 0.3% from a bolus of 3g/kg (Figure 6C) and it remained substantially stable along time. Moreover, the fraction of enhanced pixels displayed a similar percentage 47-50% for the 1.5 g/kg dose and 50-52% for the 3 g/kg dose, respectively (Figures 6B and 6D). Comparable CEST contrast enhancements and fraction of enhanced pixels were observed for the 3OMG at both the two doses. At the dose of 1.5 g/kg, we observed a ΔST% of 1.4 ± 0.2 %, whereas at the dose of 3 g/kg a ΔST% of 1.6 ± 0.2 % that remained stable after 30 min (Figure 6E and 6G). Almost similar fraction of enhanced pixels was observed among the doses: 49-53% for the 1.5 g/kg dose and 50-56% for the 3 g/kg dose, respectively (Figures 6F and 6H). Representative parametric images for both D-glucose and 3OMG at the two doses and at the several time points (6-30 min after the i.v. administration) are shown in Figure 7, depicting a clear reduction in the CEST contrast and in the coverage of the enhanced pixels inside the tumor for both the molecules at 3 T in comparison to the 7 T.

**FIGURE 6.**
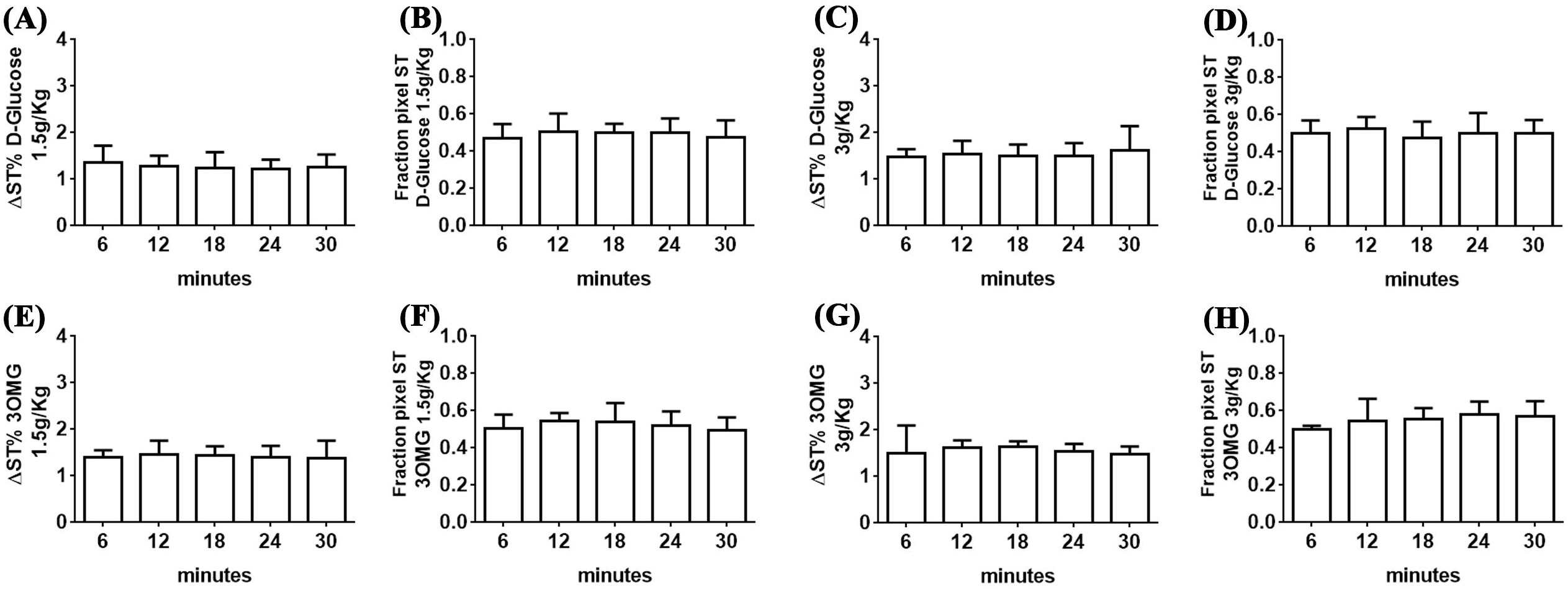
GlucoCEST contrast obtained at 3 T injecting D-Glucose at 1.5 g/kg (A) and 3 g/kg (C) dose or 3OMG at 1.5 g/kg (E) and 3 g/kg (G) dose via intravenous route and fraction of enhanced pixels at the corresponding injected dose (B, D for D-Glucose and F, H for 3OMG). Data are reported as the difference (ΔST %) between the ST effect before and after the intravenous injection.

**FIGURE 7.**
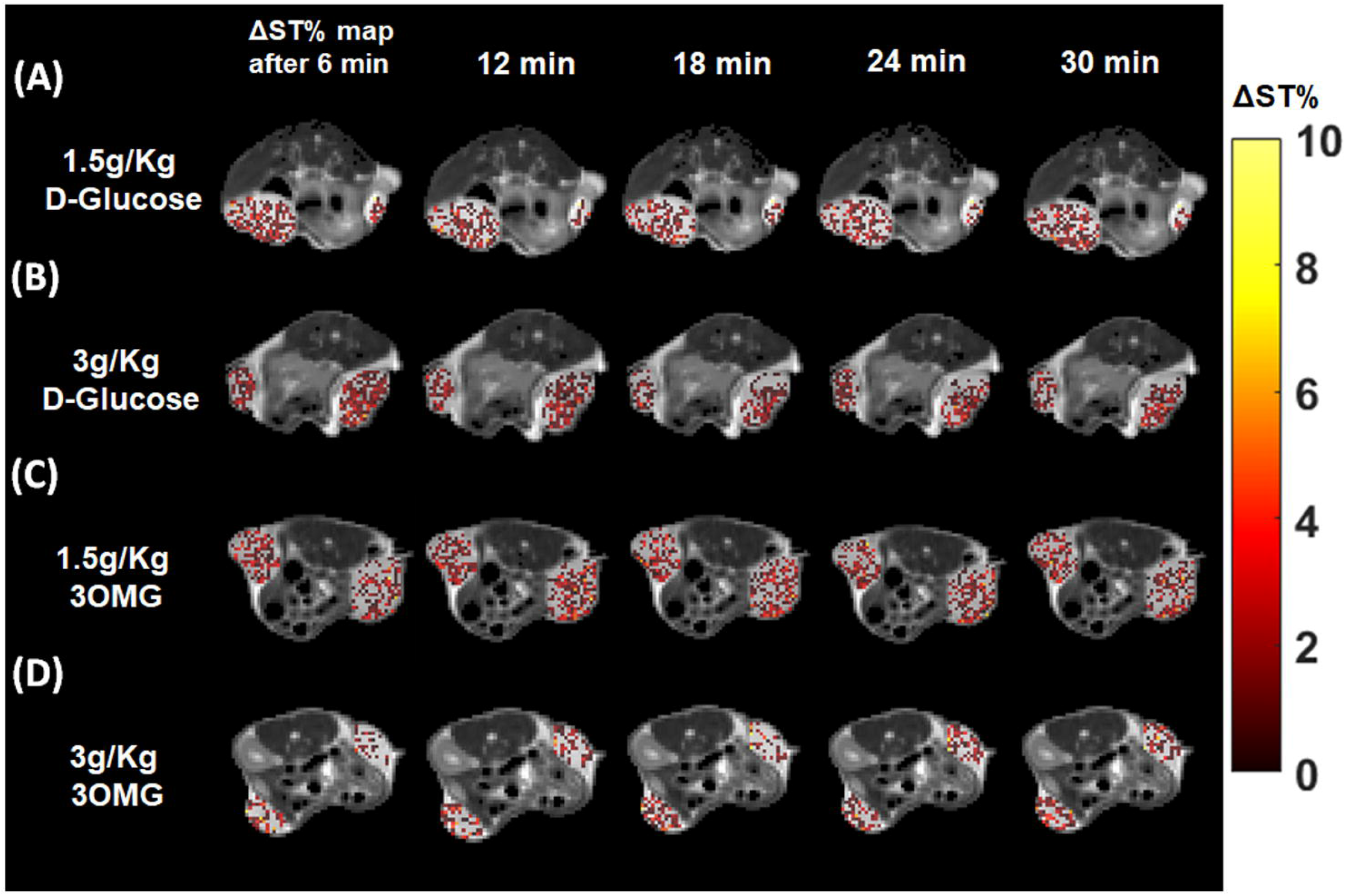
GlucoCEST ΔST% map obtained at 3 T injecting D-Glucose at 1.5 g/kg (A) and 3 g/kg (B) dose or 3OMG at 1.5 g/kg (C) and 3 g/kg (D) dose via intravenous route. Data are reported as the difference (ΔST %) between the ST effect before and after the intravenous injection, every 6 minutes from the injection. Parametric maps are overimposed to T_2w_ anatomical images and glucoCEST contrast is shown only in the tumour regions.

## 4. Discussion and Conclusions

Overall, the herein reported results highlight the role of D-Glucose and 3OMG as CEST agents for tumour detection. Both *in vitro* and *in vivo* results outline the better performance of these agents at higher magnetic field strength, due to the larger chemical shift separation between hydroxyl and water resonances. At the lower magnetic field examined, 3 T, a field strength available on clinical scanners, the separation in hertz between the hydroxyl and the bulk water protons is reduced, thus increasing the direct saturation of the bulk water pool that may decrease the labelling efficiency of the hydroxyl protons and consequently reduce the contrast performance. As expected, from higher to lower B_0_ field a decrease in CEST signals, hence in detectability, for both D-Glucose and 3OMG was observed.

The main difference between the CEST properties of D-Glucose and 3OMG that came out from the *in vitro* investigations deals with the pH dependence. In the pH range between 6 and 7.4, the extent of glucoCEST contrast for D-Glucose was higher moving towards the acidic side, whereas 3OMG displayed higher ST values when the pH was close to the neutrality, as already noted in vitro by examining the single compounds ^15,29^. Beside the opposite pH dependence between D-Glucose and 3OMG, a similar range of CEST contrasts was observed independently of the applied B_1_ power level.

In *in vivo* setting, single i.v. bolus injections of either D-Glucose or 3OMG at 7 T yielded well detectable CEST effects in the tumour region (dose 3 g/Kg, max CEST values of ca. 2-3% for D-Glucose and 3OMG, respectively). Conversely, at the magnetic field of 3 T, the average CEST effect for both contrast agents was slightly smaller but still above 1 %. Consequently, at the higher magnetic field strength of 7 T a detectable difference between the two doses was observed, whereas at lower magnetic field strength (3 T), a clear difference between the two doses was not detected. A similar decrease in the theoretically achievable glucoCEST contrast between 7 T and 3 T was recently reported from simulated data in a tumour-like tissue ^21^ and a small if not elusive CEST contrast was detected in cancer patients following glucose administration at 3T ^38,39^. Our results are in line with those reported in literature, despite the different experimental conditions (including B_0_ field, B_1_ saturation level and duration, administration route and doses, as well as image analysis) hamper any clear comparison among studies and between molecules. For examples, Van Zijl and co-workers measured a glucoCEST contrast of 3-4% after i.v. infusion of D-Glucose (dose 1 g/kg, 11.7 T) in tumour bearing mice ^15^ and at the same B_0_ field, using a dynamic glucose enhanced MRI (DGE-MRI) approach, they observed a 1% enhancement in the tumour region 5 min after the injection in a brain tumour model with a dose of 3g/kg (33). Navon and co-workers reported for the 3OMG a glucoCEST contrast of ca. 4 % 30 min after the iv injection (dose 0.7 g/kg) on implanted orthotopic mammary tumors at 7 T ^32^, whereas Van Zijl and colleagues reported, in a brain tumor model, contrast around 3 % (dose 3 g/kg on a 11.7 T) during DGE acquisition ^40^. This study provides for the first time a strict comparison in terms of the glucoCEST contrast achievable by the two investigated molecules at different magnetic fields.

Detection of CEST agents requires the accumulation in the region of interest in the order of mM of exchangeable protons concentration. Both dose regimens used in this work allowed the detection of the associated CEST effect in a large portion of the tumour region, with higher fractions of enhanced pixels when increasing the dose or the main magnetic field. As far as concerns the administration of the molecules, we investigated only the i.v. injection route since it provides the faster penetration of the agents into the tumour region and it is the common administration route for MRI contrast agents at clinical level.

Upon i.v. injection, D-glucose and 3OMG distribute in the systemic circulation and maintain high concentration levels in the tumour region. The main difference between the two agents is related to their metabolic fate: both can enter the tumour cells but only D-Glucose is metabolized, whereas 3OMG is not. On this basis, it was surmised that 3OMG, through its accumulation in the tumour cell, may act as reporter of the glucose transporters in similarly to what occurs with [^18^F]-FDG in PET assays.

The behaviour of the CEST effects detected over the 0-30 min observation time could reflect, besides variation in concentrations associated to tumor perfusion properties ^41^, changes in pH of the microenvironments in which the agents are distributed. Considering the CEST effect shown by 3OMG (Figure 4G), the observed response may arise from the extracellular contribution (that decreases upon time, characterized by an acidic pH, thus yielding a smaller ST%, as shown by the *in vitro* experiments) and the intracellular contribution (that increases upon time, characterized by a higher ST%, as shown by the in vitro experiments). Thus, the almost constancy shown over the 30 min observation time could be the balanced result of the two contributions. Vice versa, in the case of D-Glucose, the marked increase of the ST% (Figure 4C) strongly suggests a pH-drop that occurs in the extracellular matrix of the tumour cells, despite some of the molecule is metabolized inside the cancer cells. The increased glycolysis generated by the “forced” supply of glucose to the tumour cells is expected to yield, over the 30 min time, a drop in the pH of the extracellular matrix that is responsible for the observed enhanced ST%, as clearly shown in the *in vitro* results, where D-Glucose yielded an improved CEST performance at more acidic pH values. A literature-based evidence shows that the glucose challenge in tumours results in a decrease of tumour pH, following an increased glycolysis as consequence of the higher availability of glucose to the cancer cells ^42^.

Further improvements of the glucoCEST approach are ongoing. For instance, aiming at improving glucose irradiation specificity and reducing direct water saturation effects for an enhanced glucoCEST detection, the application of the chemical exchange sensitive spin-lock (CESL) technique was proposed. Jin et al. ^43^ reported robust glucoCESL signal at 9.4 T using 0.25 g/Kg of glucose. Gore and colleagues ^44^, at the same field strength, reported for the 3OMG intravenous injection (dose 1.5g/kg) an increased signal on a rat brain tumour model compared to healthy brain by appropriate analyses of MRI signals. Development of adiabatically-prepared spin-lock sequences allowed the implementation of the glucoCESL approach in clinical scanners with better visualization of glucose uptake ^22,23,35^. Recently, Zaiss and co-workers provided an accurate procedure for the optimization of the acquisition parameters in vivo based on a detailed study of the glucose -OH exchange rate at physiological conditions ^21^. Ultra high-field scanner at 17.2 T can be exploited to investigate metabolic changes induced by neuronal stimulation in rat brains with the GlucoCEST technique ^45^.

To the best of our knowledge, this is the first study in which D-Glucose and 3OMG have been systematically compared in terms of their CEST contrast efficiency at different magnetic fields and administered doses in the same murine tumour model. Further investigations are needed to compare glucose and 3OMG CEST response with the [^18^F]-FDG PET technique to assess its potential as a valid alternative on tumour diagnosis and treatment monitoring.

## Abbreviations

[^18F^]-FDG: 2-Deoxy-2-[18F]-Fluoroglucose
2DG: 2-Deoxy-D-glucose
2OMG: 2-O-Methyl-D-glucose
3OMG: 3-O-Methyl-D-glucose
6DG: 6-Deoxy-D-glucose
B0: Magnetic Field Strength
B1: Saturation Power Level
CESL: Chemical Exchange-Sensitive Spin-Lock
CEST: Chemical Exchange Saturation Transfer
CT: Computed Tomography
DGE-MRI: Dynamic Glucose Enhanced MRI
GlcN: Glucosamine
GlcNAc: N-Acetyl-Glucosamine
glucoCEST: D-Glucose Chemical Exchange Saturation Transfer
GLUT-1 and 3: Glucose Transporters 1 and 3
MRI: Magnetic Resonance Imaging
PBS: Phosphate-Buffered Saline
PET: Positron Emission Tomography
ROI: Region of Interest
ST: Saturation Transfer

## ACKNOWLEDGMENTS

We acknowledge the support received from the European Union’s Horizon 2020 research and innovation programme (Grant Agreement No. 667510), from the Associazione Italiana Ricerca Cancro (AIRC MFAG 2017 - ID. 20153 project) and from Compagnia San Paolo project (Regione Piemonte, grant #CSTO165925). MC was supported by fellowship from the Associazione Italiana Ricerca Cancro (AIRC ID 24104). The Italian Ministry for Education and Research (MIUR) is gratefully acknowledged for yearly FOE funding to the Euro-BioImaging Multi-Modal Molecular Imaging Italian Node (MMMI).

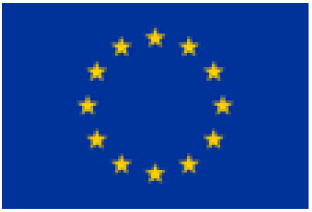

## DATA AVAILABILITY

The data that support the findings of this study are available from the corresponding author upon reasonable request.

